# The COMA complex is required for positioning Ipl1 activity proximal to Cse4 nucleosomes in budding yeast

**DOI:** 10.1101/444570

**Authors:** Josef Fischböck Halwachs, Sylvia Singh, Mia Potocnjak, Götz Hagemann, Victor Solis, Stephan Woike, Medini Ghodgaonkar Steger, Jessica Andreani Feuillet, Franz Herzog

## Abstract

Kinetochores are macromolecular protein complexes assembled on centromeric chromatin that ensure accurate chromosome segregation by linking DNA to spindle microtubules and integrating safeguard mechanisms. A kinetochore-associated pool of Ipl1^Aurora B^ kinase, a subunit of the chromosomal passenger complex (CPC), was previously implicated in feedback control mechanisms. To study the kinetochore subunit connectivity built on budding yeast point centromeres and its CPC interactions we performed crosslink-guided *in vitro* reconstitution. The Ame1/Okp1^CENP-U/Q^ heterodimer, forming the COMA complex with Ctf19/Mcm21^CENP-P/O^, selectively bound Cse4^CENP-A^ nucleosomes through the Cse4 N-terminus and thereby establishes a direct link to the outer kinetochore MTW1 complex. The Sli15/Ipl1^INCENP/Aurora B^ core-CPC interacted with COMA through the Ctf19 C-terminus, and artificial tethering of Sli15 to Ame1/Okp1 rescued synthetic lethality upon Ctf19/Mcm21 deletion in a Sli15 centromere-targeting deficient mutant. This study reveals characteristics of the inner kinetochore architecture assembled at point centromeres and the relevance of its Sli15/Ipl1 interaction for CPC function.

## Introduction

Kinetochores enable the precise distribution of chromosomes during the eukaryotic cell division to avoid aneuploidy (Santaguida and Musacchio, 2009) which is associated with tumorigenesis, congenital trisomies and aging (Baker et al., 2005, Pfau and Amon, 2012). Faithful segregation of the duplicated sister chromatids relies on their exclusive attachment to spindle microtubules emerging from opposite spindle poles (Foley and Kapoor, 2013). The physical link between chromosomal DNA and microtubules is formed by the kinetochore, a macromolecular protein complex, which mediates the processive binding to depolymerizing microtubules driving the sister chromatids apart into the two emerging cells (Biggins, 2013, Musacchio and Desai, 2017). Kinetochore assembly is restricted to centromeres, chromosomal domains that are epigenetically marked by the presence of the histone H3 variant Cse4^CENP-A^ (human ortholog names are superscripted if appropriate) (Earnshaw and Rothfield, 1985, Fukagawa and Earnshaw, 2014). In humans, regional centromeres span megabases of DNA embedding up to 200 CENP-A containing nucleosomal core particles (NCPs) (Bodor et al., 2014, Musacchio and Desai, 2017). In contrast, *Saccharomyces cerevisiae* possesses point centromeres which are characterized by a specific ~125 bp DNA sequence wrapped around a single Cse4 containing histone octamer. The kinetochore unit built on this point centromere stays assembled and bound to a single microtubule throughout almost the whole cell cycle (Pluta et al., 1995, Biggins, 2013, Lang et al., 2018).

The budding yeast kinetochore is composed of about 45 core subunits which are organized in different stable complexes (De Wulf et al., 2003, Westermann et al., 2003) of which several are present in multiple copies (Joglekar et al., 2006). The kinetochore proteins are evolutionary largely conserved between yeast and humans (Westermann and Schleiffer, 2013, van Hooff et al., 2017) and share a highly similar hierarchy of assembly from DNA to the microtubule binding interface (Santaguida and Musacchio, 2009) The centromere proximal region is established by proteins of the Constitutive Centromere Associated Network (CCAN), also referred to as Ctf19 complex in budding yeast, which is assembled by Mif2^CENP-C^ and the Chl4/Iml3^CENP-N/L^, Mcm16/Ctf3/Mcm22^CENP-H/I/K^, Cnn1/Wip1^CENP-T/W^, Mhf1/Mhf2^CENP-S/X^, Ctf19/Okp1/Mcm21/Ame1^CENP-P/Q/O/U^ (COMA) complexes (Musacchio and Desai, 2017) including the budding yeast specific yeast Nkp1/Nkp2 and CBF3 (Ndc10/Cep3/Ctf13/Skp1) complexes (Lechner and Carbon, 1991, Cho and Harrison, 2011, Cheeseman et al., 2002). The CCAN complex provides a cooperative high-affinity binding environment for the Cse4^CENP-A^-NCP (Weir et al., 2016) where distinct subunits selectively recognize Cse4^CENP-A^ specific features. Across different species the CENP-C signature motif interacts with divergent hydrophobic residues of the CENP-A C-terminal tail (Musacchio and Desai, 2017). Electron microscopy studies have recently resolved the interaction of CENP-N with the CENP-A centromere-targeting domain (CATD) of CENP-A-NCP in vertebrates (Carroll et al., 2009, Guse et al., 2011, Pentakota et al., 2017, Chittori et al., 2018, Tian et al., 2018). For budding yeast Cse4 a direct interaction has so far only been demonstrated with Mif2 (Westermann et al., 2003, Xiao et al., 2017). Interestingly, apart from Mif2 the only essential CCAN proteins are Ame1 and Okp1 (Meluh and Koshland, 1997, Ortiz et al., 1999, De Wulf et al., 2003), with the N-terminus of Ame1 binding the N-terminal domain of Mtw1 and serving as docking site for the outer kinetochore KMN network (KNL1^Spc105^-/MIS12^MTW1^-/NDC80^Ndc80^-complexes) (Hornung et al., 2014, Dimitrova et al., 2016).

The kinetochore is also a hub for feedback control mechanisms that ensure high fidelity of sister chromatid separation by aligning the microtubule attachment state to cell cycle progression, known as spindle assembly checkpoint (SAC), and by selectively destabilizing improper kinetochore-microtubule attachments and stabilizing the correct bipolar attachment, referred to as error correction mechanism (Foley and Kapoor, 2013, Krenn and Musacchio, 2015). A major effector of both regulatory feedback loops is the kinase Ipl1^Aurora B^, a subunit of the evolutionary conserved tetrameric Chromosomal Passenger Complex (CPC), which associates close to the centromere from G1 until anaphase (Biggins and Murray, 2001, Widlund et al., 2006, Carmena et al., 2012). The CPC is built-up by Ipl1^Aurora B^ which binds to the C-terminal IN-box domain (Adams et al., 2000, Kaitna et al., 2000) of the scaffold protein Sli15^INCENP^, whereas Nbl1^Borealin^ and Bir1^Survivin^ form a three-helix bundle with the Sli15 N-terminus (Klein et al., 2006, Jeyaprakash et al., 2007). All known mechanisms for recruitment of the CPC to the budding yeast centromere rely on Bir1 which directly associates with Ndc10 of the CBF3 complex and is recruited through Sgo1 to histone H2A phosphorylated by Bub1 (Kawashima et al., 2010, Cho and Harrison, 2011). During early mitosis incorrect microtubule attachment states are resolved by Ipl1^AuroraB^ which phosphorylates Ndc80 and Dam1 sites within the microtubule binding interface and thereby reduces the affinity towards microtubules (Cheeseman et al., 2002, Cheeseman et al., 2006, DeLuca et al., 2006, Westermann et al., 2005, Miranda et al., 2005, Santaguida and Musacchio, 2009). The selective destabilization promotes the establishment of a correctly bi-oriented kinetochore configuration at the mitotic spindle, referred to as amphitelic attachment (Liu et al., 2009). One model for establishing biorientation (Krenn and Musacchio, 2015) implies that centromere-targeting of Sli15 allows substrate phosphorylation by Ipl1^Aurora B^ within the span of the Sli15^INCENP^ scaffold and that tension dependent intra-kinetochore stretching pulls the microtubule binding interface out of reach of Ipl1^Aurora B^ resulting in dephosphorylation and stabilization (Lampson and Cheeseman, 2011). A recent study by Campbell & Desai challenged this model by showing that a mutant version of Sli15 lacking the centromere-targeting domain Sli15∆N2-228 (Sli15∆N), still colocalizes with kinetochores, is indistinguishably viable from wild-type and displays no significant chromosome segregation defects (Campbell and Desai, 2013). Similarly, a survivin mutant in chicken DT40 cells that fails to localize INCENP and Aurora B to centromeres from prophase to metaphase has normal growth kinetics (Yue et al., 2008). These findings suggest that centromere-targeting of Sli15/Ipl1 is largely dispensable for error correction and SAC signaling. The molecular mechanism of how centromere-targeting deficient Sli15/Ipl1 is recruited to kinetochores is unclear.

Here, we use chemical crosslinking and mass spectrometry (XLMS) (Herzog et al., 2012) together with biochemical reconstitution to characterize the CCAN subunit topology and the binding interfaces that establish a selective Cse4-NCP binding environment. Interestingly, subunits of the COMA complex were previously implicated in CPC function at kinetochores (De Wulf et al., 2003, Knockleby and Vogel, 2009) and the centromere-targeting deficient Sli15ΔN mutant shows synthetic lethality with deletion of Ctf19/Mcm21 (Campbell and Desai, 2013). Thus, we investigated whether the COMA complex directly associates with Sli15/Ipl1 and whether these interactions are relevant *in vivo*. We demonstrate that the Cse4-N-terminus (Chen et al., 2000) recruits the Ame1/Okp1 heterodimer through the Okp1 core domain (Schmitzberger et al., 2017) and that dual recognition of budding yeast Cse4-NCP is established through selective interactions of the essential CCAN proteins Mif2 and Ame1/Okp1 with distinct Cse4 motifs. We further reveal that Sli15/Ipl1 interacts with the Ctf19 C-terminus and that synthetic lethality upon depletion of Ctf19 in the Sli15 centromere-targeting deficient *sli15ΔN* background is rescued exclusively by fusing Sli15ΔN to the COMA complex. Together these findings reveal a distinct CCAN architecture assembled at budding yeast point centromeres and indicate that the interaction of CPC and COMA complexes may position Ipl1 close to Cse4 nucleosomes in order to mediate feedback control.

## Results

### The Ame1/Okp1 heterodimer selectively binds Cse4 containing nucleosomes

Previous *in vitro* reconstitution experiments using mammalian proteins highlighted a specific hierarchical assembly of CCAN complexes at nucleosomes with CENP-C and CENP-N recognizing CENP-A (Carroll et al., 2009, Carroll et al., 2010, Kato et al., 2013, Klare et al., 2015, Weir et al., 2016, Guse et al., 2011, Falk et al., 2016). Due to the spatially restricted point centromere in budding yeast, cooperative stable interactions between the single Cse4 nucleosome and kinetochore proteins are anticipated. Up to now, only Mif2 was shown to directly interact with Cse4 (Xiao et al., 2017). To screen for further direct interaction partners of Cse4-NCPs we reconstituted the individual CCAN subcomplexes (Mif2, Ame1/Okp1, Ctf19/Mcm21, Chl4/Iml3, Mcm16/Ctf3/Mcm22, Cnn1/Wip1, Nkp1/Nkp2, Mhf1/Mhf2) with Cse4- or H3-NCPs *in vitro*. The CCAN complexes were purified either from bacteria or insect cells as homogenous and nearly stoichiometric complexes (Figure 1B). Consistent with a recent study (Xiao et al., 2017) using electrophoretic mobility shift assays (EMSA), we observed that Mif2 selectively interacted with Cse4-NCPs and not with H3-NCPs (Figure 1A). Notably, we identified Ame1/Okp1 to form a complex with Cse4-NCPs (Figure 1A). The lack of interaction with H3-NCPs, which was reconstituted using the same 601 DNA sequence (Tachiwana et al., 2011), suggests that Ame1/Okp1 directly and selectively binds Cse4 and not AT-rich DNA sequences as previously proposed (Hornung et al., 2014). In contrast to findings in humans, Chl4/Iml3, the orthologs of human CENP-N/L (Carroll et al., 2009), did not shift the Cse4-NCP band in the EMSA. Similarly, EMSA band shifts observed for human CENP-A-NCPs with the CENP-HIK complex (Weir et al., 2016) could not be reproduced with the budding yeast orthologs. None of the other Ctf19 subcomplexes, Ctf19/Mcm21, Cnn1/Wip1, Nkp1/Nkp2 and Mhf1/Mhf2, associated with Cse4- or H3-NCPs in the EMSA (Figure 1A).

**Figure 1.**
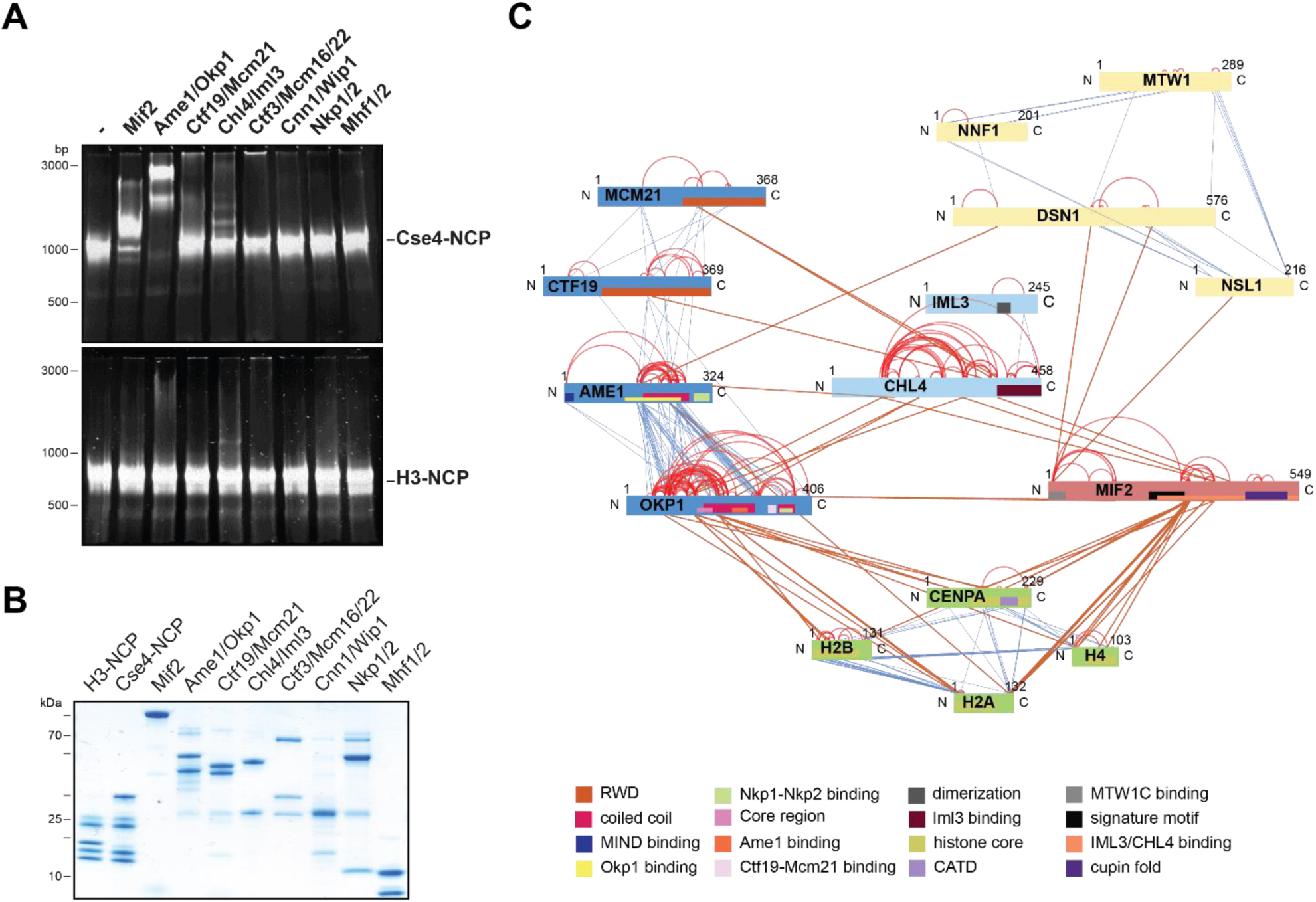
The heterodimeric Ame1/Okp1 complex directly and selectively binds the Cse4-NCP. (**A**) Electrophoretic mobility shift assays (EMSAs) of the indicated CCAN subunits and subcomplexes mixed with either Cse4- or H3-NCPs. DNA/protein complexes were separated on a 6% native polyacrylamide gel. The DNA is visualised by fluorescent staining with SYBR Gold. (**B**) Coomassie stained gel of the individual proteins used in the EMSA in (A). (**C**) XLMS analysis of the *in vitro* reconstituted Cse4-NCP:Mif2:COMA:Chl4/Iml3:MTW1c complex. Proteins are represented as bars indicating annotated domains according to the color scheme in the legend. Subunits of a complex are represented in the same color and protein lengths and crosslink sites are scaled to the amino acid sequence.

To elucidate the binding interfaces of the Ame1/Okp1:Cse4-NCP complex we performed XLMS analysis. In order to gain insights into the protein network linked to Cse4-NCPs we reconstituted a complex composed of Ame1/Okp1, Mif2, Ctf19/Mcm21, Chl4/Iml3 and the MTW1c which links the KMN network to the inner kinetochore receptors Ame1 and Mif2 (Screpanti et al., 2011, Przewloka et al., 2011, Hornung et al., 2014). In total 349 inter-subunit crosslinks between the fifteen proteins were identified (Figure 1C, Table S3). The majority of the crosslinks detected within the different subcomplexes MTW1c, COMA, Chl4/Iml3, and Cse4-NCP are in agreement with previous studies validating our crosslink map (Hornung et al., 2014, De Wulf et al., 2003, Hinshaw and Harrison, 2013). Moreover, the crosslinks from Mif2 to the MTW1c (Screpanti et al., 2011, Przewloka et al., 2011), to Chl4 (Hinshaw and Harrison, 2013) and to Cse4 (Kato et al., 2013) are consistent with previously described interfaces. In particular, the distance restraints which connect the Mif2 N-terminus and MTW1c, link the Mif2 Chl4/Iml3 binding domain to Chl4 and position the Mif2 signature motif next to the Cse4 C-terminus (Figure 1C, Table S3). Interestingly, crosslinks between Ame1/Okp1 and Cse4 occur exclusively between Okp1 and Cse4 suggesting that Okp1 is the direct binding partner of Cse4. Furthermore, Okp1 was the only COMA subunit that crosslinked to the three canonical histones H2A, H2B and H4 with the exception of one crosslink between Ame1 and H2A. Notably, our analysis revealed crosslinks between Chl4 and all COMA subunits suggesting a direct association between Chl4/Iml3 and COMA complexes. The human orthologs CENP-OPQUR were recently shown to bind through a composite interface to CENP-LN and CENP-HIKM (Pesenti et al., 2018), however, no direct interaction between COMA and Chl4/Iml3 complexes has been reported. Taken together, the EMSA and XLMS analyses identified a previously unknown interaction between Cse4 and the Ame1/Okp1 heterodimer which is putatively mediated through Okp1 and revealed that Chl4 does not recognize Cse4-NCPs in contrast to its human ortholog.

### The essential N-terminal domain of Cse4 is required for Okp1 binding

To further characterize the interaction between Ame1/Okp1 and Cse4-NCPs we aimed to identify the binding interface of the Ame1/Okp1:Cse4-NCP complex. Importantly, two crosslinks were detected between Okp1 and the essential Cse4 N-terminus (Figure 1C, Table S3). A multiple sequence alignment (MSA) of Cse4^CENP-A^ protein sequences (Figure 2A) unveiled a conserved region (ScCse4 aa 34-61) unique to Cse4 proteins of inter-related yeasts which is almost identical to the essential N-terminal Domain (END, aa 28-60). The END domain was previously shown to be required for Cse4 function and for recruiting the ‘Mcm21p/Ctf19p/Okp1p complex’ to minichromosomes (Chen et al., 2000, Ortiz et al., 1999). A molecular understanding of the mechanistic role of the essential END domain is still missing.

**Figure 2.**
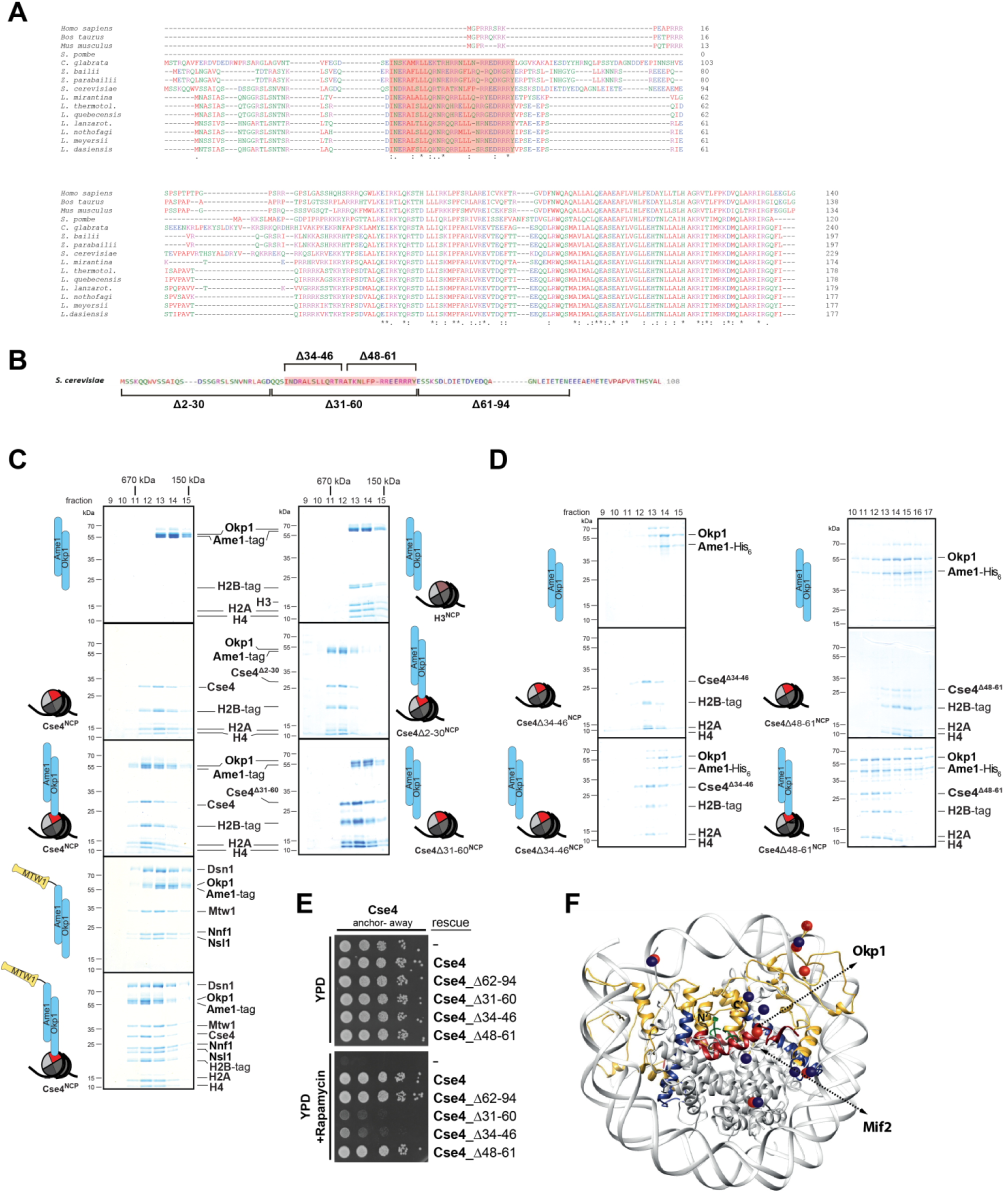
A short helical motif within the Cse4 N-terminus serves as Ame1/Okp1 docking site and is essential *in vivo*. (**A**) Multiple sequence alignment of Cse4^CENP-A^ proteins. Yeast protein sequences with the highest similarities to *S. cerevisiae* Cse4, three mammalian and the *S. pombe* homologous CENP-A protein sequences were included in the alignment. The amino acid (aa) patch, conserved in interrelated yeasts, is highlighted in pink (*S. cerevisiae* Cse4 aa 34-61). Amino acid residues are colored and annotated according to the ClustalW color and annotation codes (S.: *Schizosaccharomyces*, C.: *Candida*, Z.: *Zygosaccharomyces*, L.: *Lachancea*). (**B**) Scheme of the deletion mutants within the Cse4 N-terminus used in the SEC experiments in (C) and (D) and in the cell viability assays in (E). The conserved region (aa 34-61) is highlighted in pink. (**C**) SEC analysis of the indicated mixtures of recombinant Ame1/Okp1 and MTW1c and reconstituted H3-, Cse4-, Cse4Δ2-30- or Cse4Δ31-60-NCPs. Ame1/Okp1, MTW1c and the Cse4 proteins were mixed equimolar. Eluted proteins were visualized by SDS-PAGE and Coomassie staining. (**D**) SEC analysis of Ame1/Okp1 preincubated with Cse4Δ34-46- or Cse4Δ48-61-NCPs. Eluted complexes were analyzed by SDS-PAGE and Coomassie staining. (**E**) Cell growth assay of Cse4 mutants in budding yeast using the anchor-away system. The Cse4 wild-type and indicated mutant proteins were ectopically expressed in a Cse4 anchor-away strain (Cse4-FRB) and cell growth was monitored by plating 1:10 serial dilutions on YPD medium at 30 °C in the absence or presence of rapamycin. (**F**) Structural model of the *S. cerevisiae* Cse4-NCP. The Cse4 histone is highlighted in gold. The histone-fold domain of Cse4 harbors the CATD (blue) and the C-terminus (dark green) which is essential for the recruitment of Mif2. The N-terminal tail of Cse4 carries the END domain (dark red) which mediates binding to Okp1 (dark red). Cse4 residues crosslinked to Mif2 or Okp1 are indicated by dark blue or red spheres, respectively. The selective dual recognition of Cse4-NCP is achieved by Mif2 and Ame1/Okp1 interacting with distinct sequence motifs of the Cse4 protein.

To assess whether the Cse4 END domain mediates interaction with the Ame1/Okp1 heterodimer we tested binding of recombinant Ame1/Okp1 to reconstituted wild-type and deletion mutants (Figure 2B) of Cse4- and H3-NCPs using size exclusion chromatography (SEC) (Figure 2C). Wild-type Cse4-NCP but not H3-NCP formed a stoichiometric complex with Ame1/Okp1 (Figure 2C) which is consistent with our findings of the EMSA and XLMS analyses (Figure 1 A, C). In addition, Ame1/Okp1 bound to a Cse4-NCP retained the ability to interact with the MTW1c (Hornung et al., 2014) forming a supramolecular complex with Ame1 and Okp1 directly linking the KMN network to the centromeric nucleosome (Figure 2C). Truncation of the first 30 N-terminal residues of Cse4 neither affected its ability to bind Ame1/Okp1 nor was it essential for viability (Figure 2C) (Chen et al., 2000). However, deletion of the END domain in Cse4∆31-60 abrogated Ame1/Okp1:Cse4-NCP complex formation (Figure 2C). To further narrow down the interface, two deletion mutants splitting the END domain in half, Cse4∆34-46 and Cse4∆48-61 (Figure 2B), were tested in SEC experiments. While Cse4∆48-61 associated with Ame1/Okp1, deletion of amino acids 34-46 completely disrupted the interaction (Figure 2D). All Cse4 N-terminal mutants and wild-type NCPs eluted at similar retention times from SEC indicating that the Cse4 N-terminal deletion mutants did not affect Cse4 incorporation and stability of the nucleosomes.

The crosslink-derived distance restraints as well as testing complex formation of wild-type and mutant Cse4 proteins by SEC identified a conserved Cse4 peptide motif of amino acids 34-46 to be sufficient for Ame1/Okp1 interaction. To test whether this motif is also essential for cell viability we depleted endogenous Cse4 from the nucleus using the anchor-away technique and performed rescue experiments by ectopically expressing the Cse4 mutants Cse4∆34-46 and Cse4∆48-61. Indeed, deletion of amino acids 34-46 was lethal whereas the mutant lacking residues 48-61 displayed wild-type growth rates (Figure 2E). The observation that deletion of the minimal Ame1/Okp1 interacting Cse4 motif (aa 34-46) correlates with the loss of cell viability whereas the C-terminal half of the END domain (aa 48-61) is neither essential for viability nor required for Ame1/Okp1 association suggests that recruitment of the Ame1/Okp1 heterodimer to Cse4-NCPs through the minimal Ame1/Okp1 interacting Cse4 motif is essential for yeast growth. The Mif2 signature motif (Xiao et al., 2017) and Ame1/Okp1 recognize distinct motifs at the Cse4 C- and N-terminus (Figure 2F), respectively, and both interactions are essential for viability (Hornung et al., 2014).

### The Okp1 core domain interacts with Cse4

To characterize the Cse4 binding site in Okp1 we applied crosslink-derived restraints to narrow down the putative interface to amino acids 95-202 of Okp1 (Figure 1C, Table S3). Based on MSA analysis of Okp1 sequences, this region harbors a conserved stretch (aa 127-184), including part of the previously described Okp1 core domain (aa 166-211) which is essential for cell growth and whose function is still elusive (Schmitzberger et al., 2017) (Figure 3A). Furthermore, a secondary structure analysis predicted two alpha helices within the conserved domain (helix1 aa 130-140, helix2 aa 156-188) (Figure 3A). Thus, we designed three deletion mutants (Okp1Δ123-147, Okp1Δ140-170, Okp1Δ163-187) and purified all Okp1 mutant proteins in complex with Ame1 from *E*. *coli*. In EMSAs Ame1/Okp1Δ123-147 bound to Cse4-NCPs similarly to the wild-type Ame1/Okp1 complex whereas Ame1/Okp1Δ140-170 only weakly associated and Ame1/Okp1Δ163-187 did not form a complex with Cse4-NCP (Figure 3B). These results are consistent with monitoring protein complex formation using SEC (Figure S1). In addition, analysis of the Okp1 deletion mutants Δ123-147 and Δ163-187 in cell viability assays again revealed a tight correlation between their requirement for the interaction with Cse4 and being essential for yeast growth (Figure 3C) (Schmitzberger et al., 2017). This finding further supports the notion that the recognition of the Cse4 nucleosome by Ame1/Okp1 is essential in budding yeast.

**Figure 3.**
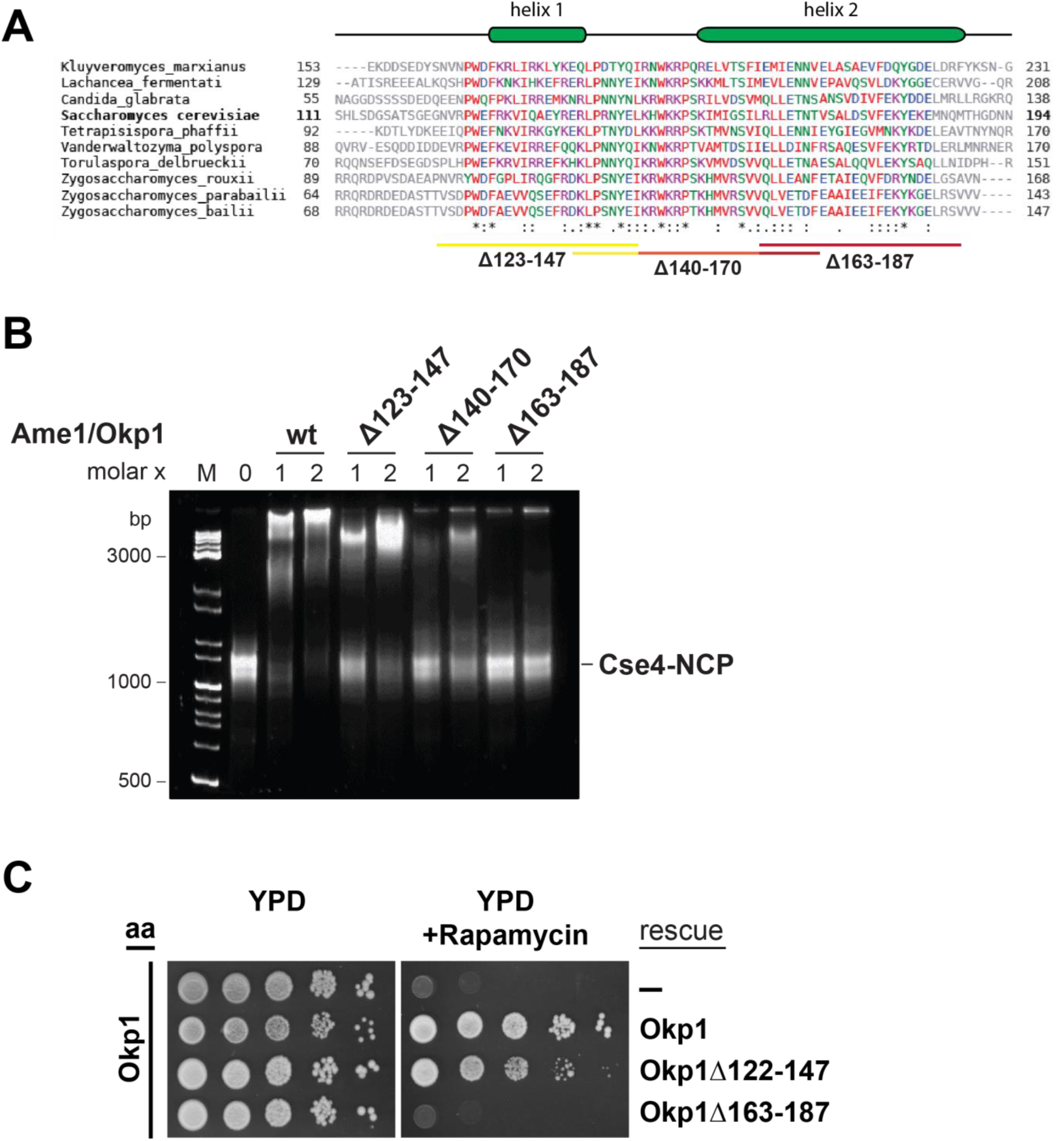
The essential core domain of Okp1 is required for the interaction with Cse4-NCPs. (**A**) Multiple sequence alignment of Okp1 amino acid sequences from related yeast species. Amino acid residues of the conserved region are colored and annotated according to the ClustalW color and annotation codes. Green bars above the alignment represent alpha helical regions predicted by Jpred (Drozdetskiy et al., 2015). Lines below the alignment indicate the overlapping Okp1 deletion mutants analyzed in (B) and (C). (**B**) EMSA monitoring complex formation of Cse4-NCPs with Ame1/Okp1 wild-type (wt) and heterodimers including Okp1 deletion mutants. Recombinant Ame1/Okp1 complexes were tested in a 1:1 and 2:1 molar ratio to Cse4-NCPs. The DNA is visualised by fluorescent staining with SYBR Gold. (**C**) Cell viability assay of Okp1 deletion mutants using the anchor away technique. Yeast growth of either the untransformed (-) Okp1 anchor-away strain (Okp1-FRB) or of strains transformed with the indicated Okp1 rescue alleles was tested in 1:10 serial dilutions on YPD medium in the absence or presence of rapamycin for 72 hours at 30 °C.

### The COMA complex interacts with Sli15/Ipl1 through the Ctf19 C-terminus

The COMA complex is composed of two essential, Ame1/Okp1, and two non-essential, Ctf19/Mcm21, subunits (Ortiz et al., 1999, Cheeseman et al., 2002). Both, Ctf19 and Mcm21 contain C-terminal tandem-RWD (RING finger and WD repeat containing proteins and DEAD-like helicases) domains forming a rigid heterodimeric Y-shaped scaffold whose respective N-terminal RWDs of the tandems pack together as revealed by a recent crystal structure of the *K. lactis* complex (Schmitzberger and Harrison, 2012). As recently shown, *ctf19Δ* or *mcm21Δ* mutants become synthetically lethal in a Sli15 centromere-targeting deficient background *(sli15ΔN*) (Campbell and Desai, 2013). Furthermore, Ame1 was suggested to have a role in Sli15 localization close to kinetochores independently of Bir1 (Knockleby and Vogel, 2009). To investigate whether Sli15/Ipl1 associates with the COMA complex, *in vitro* reconstitution and XLMS analysis detected 98 inter-protein and 69 intra-protein crosslinks (Figure 4A, Table S4). Notably, 10 crosslinks were found from the C-terminal RWD (RWD-C) domain of Ctf19 and 4 crosslinks from the RWD-C domain of Mcm21 to the microtubule binding domain of Sli15 (aa 229-565) (Figure 4A, Table S4, S5). Within the Ame1/Okp1 heterodimer 2 crosslinks from Sli15 to Okp1 and 1 crosslink from Ipl1 to Ame1 were identified. Interestingly, the crosslink detected to lysine 366 of Okp1 is located near the identified Ctf19/Mcm21 binding site within Okp1 (‘segment 1’ aa 321-329) (Schmitzberger et al., 2017) and thus is in close proximity to the RWD-C domains of Ctf19 and Mcm21. The interaction of Sli15 to the Ctf19 RWD-C domain indicated by crosslinks was verified in *in vitro* binding assays. Sli15-2xStrep/Ipl1 was immobilised on Streptavidin beads and incubated with 2-fold molar excess of either Ame1/Okp1 or Ctf19/Mcm21 or both using wild-type Ctf19 protein or a C-terminal deletion mutant Ctf19Δ270-369 (Ctf19ΔC). Ame1/Okp1 and Ctf19/Mcm21 were both pulled down with Sli15/Ipl1 either as individual complexes or in combination. In agreement with previous findings (Schmitzberger et al., 2017) recombinant Ctf19ΔC formed a stoichiometric complex with Mcm21, however, completely lost its ability to bind Sli15/Ipl1 validating the RWD-C of Ctf19 as primary Sli15/Ipl1 interaction site within Ctf19/Mcm21 *in vitro* (Figure 4B). Notably, autophosphorylation of Sli15/Ipl1 *in vitro* abrogated its interaction with Ame1/Okp1 and Ctf19/Mcm21 indicating that similar to microtubule binding, autophosphorylation of Sli15 by Ipl1 may prevent and regulate its binding to the COMA complex (Figure 4B, Figure S2).

**Figure 4.**
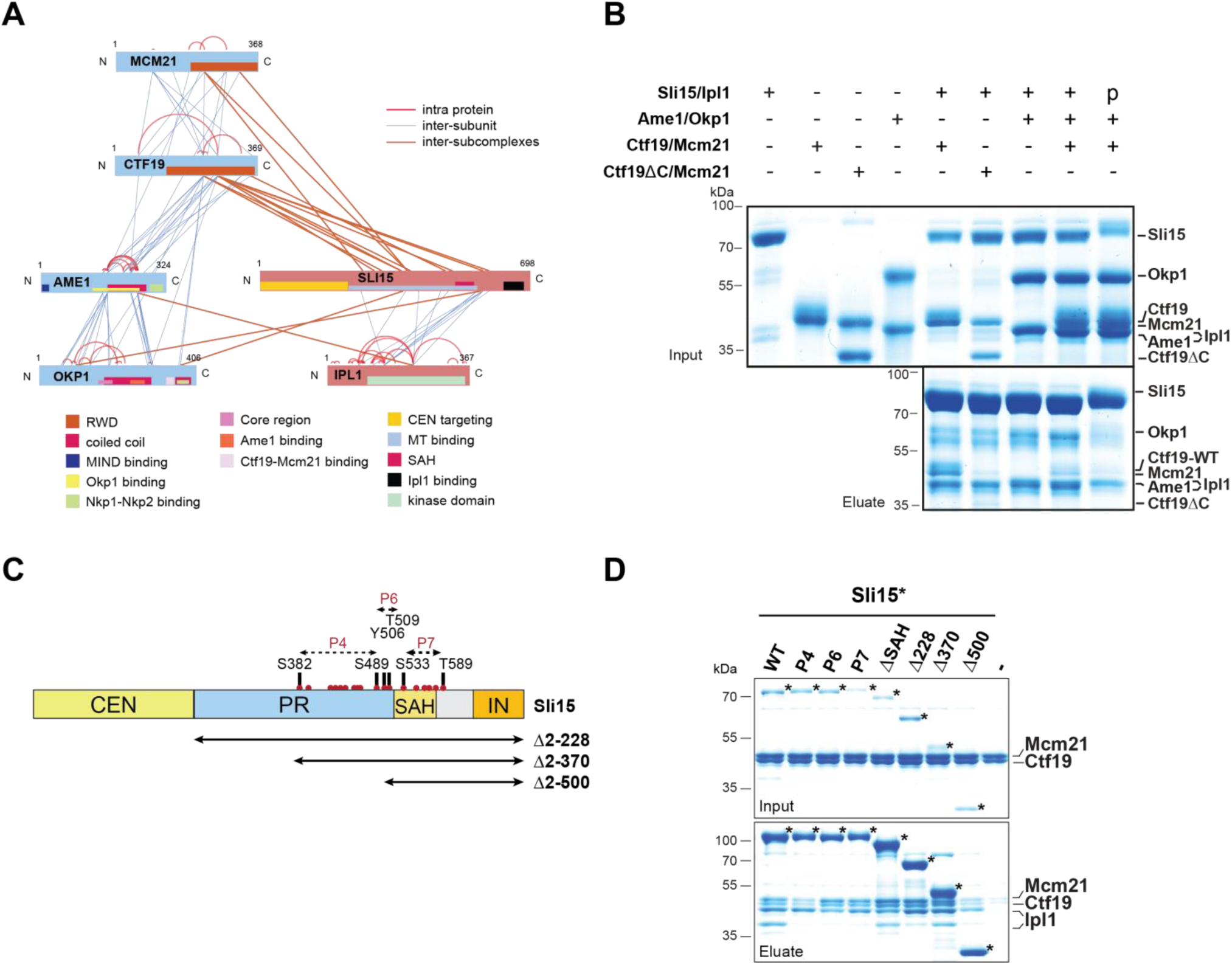
The core CPC Sli15/Ipl1 associates with the COMA complex through the Ctf19 C-terminal RWD domain *in vitro*. (**A**) Network representation of lysine-lysine crosslinks identified on recombinant Sli15/Ipl1 in complex with COMA. Proteins are represented as bars indicating annotated domains according to the color scheme in the legend. Subunits of a complex are represented in the same color. Protein lengths and crosslink sites are scaled to the amino acid sequence. (**B**) *In vitro* binding assay analyzing the interaction of Sli15/Ipl1 with the COMA complex. Recombinant Sli15-2xStrep/Ipl1 was immobilized on Streptavidin beads and incubated with Ctf19/Mcm21, Ctf19∆C/Mcm21, Ame1/Okp1 or Ame1/Okp1/Ctf19/Mcm21. Ctf19∆C lacks the last 100 amino acids which form the C-terminal RWD domain. Autophosphorylation (p) of Sli15/Ipl1 largely reduced bound protein levels which were visualized by SDS-PAGE and Coomassie staining. (**C**) Scheme depicting Sli15 wild-type and the deletion and phosphorylation site mutants tested in (D). Centromere targeting domain (CEN, aa 2-228) in yellow; phospho-regulated region (PR, aa 229-515) in light blue; single alpha helix (SAH, aa 516-575) in dark yellow; Ipl1 binding IN-box (IN, aa 626-698) in orange. The Sli15 N-terminal deletion mutants are indicated by arrows below the scheme. The patches spanning the phospho-mimetic mutants P4 (S382D, S383D, S425D, S427D, T429D, S437D, T457D, S460D, S462D, S489D), P6 (Y506E, T509D) and P7 (S533D, 554D, T564D, S567D, S578D, T589D) are indicated by arrows above the scheme. The positions of the individual phosphorylation sites are labeled by red dots. (D) *In vitro* binding assay testing the interaction of Sli15 phospho-mimicking and deletion mutants with Ctf19/Mcm21. Sli15-2xStrep/Ipl1 was immobilized on a Streptavidin matrix and eluted proteins were visualized by SDS-PAGE and Coomassie staining. The positions of the Sli15 wild-type and mutant proteins are indicated by stars.

Since crosslinks between Ctf19 and Sli15 were detected in the Sli15 microtubule binding domain (Figure 4A, Table S4, Table S5) we identified the autophosphorylation sites by mass spectrometry (Figure 4C and S2) and tested various Sli15 phosphorylation-mimicking and deletion mutants for their ability to disrupt binding to Ctf19/Mcm21 *in vitro* (Figure 4D). Sli15 wild-type and mutant proteins were immobilized on beads and two Sli15 mutants retained significantly reduced protein levels of wild-type Ctf19/Mcm21. Sli15∆N2-370 bound Ctf19/Mcm21 whereas Sli15∆N2-500 lost its ability to interact with Ctf19/Mcm21, similarly to the Sli15-P4 mutant, which carries 10 phosphorylation-mimicking mutations in the region of aa 380-490. These results indicate that Ctf19/Mcm21 associates with the Sli15 region of aa 370-500 and that Sli15/Ipl1 autophosphorylation in this region may negatively regulate this interaction.

Taken together, crosslink-derived restraints pinpointed the Ctf19 RWD-C domain as Sli15/Ipl1 docking site within the COMA complex which was supported by the loss of interaction upon deletion of the Ctf19 C-terminus *in vitro*.

### Tethering Sli15ΔN selectively to the COMA complex rescues the synthetic lethality of *sli15ΔN*/*ctf19Δ* double mutants

Intriguingly, *Ctf19* or *Mcm21* deletion is synthetically lethal with the centromere-targeting deficient *sli15ΔN* mutant, but the underlying molecular mechanism is still unknown. Since Ctf19 also has a role in centromeric cohesin loading (Fernius and Marston, 2009), the synthetic lethality of *sli15ΔN* has been attributed to a defect in cohesion establishment or maintenance (Campbell and Desai, 2013). To understand the mechanims of the synthetic effect we performed various yeast viability assays.

First, we reproduced the reported synthetic lethality using the anchor-away system (Haruki et al., 2008), where the endogenous Ctf19-FRB is inducibly depleted from the nucleus upon addition of rapamycin to the medium. CTF19-FRB depletion was performed in a yeast strain, where the endogenous *Sli15* copy was replaced by *sli15ΔN*. We found that in the presence of Ctf19-FRB, cells expressing Sli15ΔN are viable and display synthetic lethality upon Ctf19-FRB depletion consistent with previous findings (Campbell and Desai, 2013) (Figure 5A). The recent identification of the Ctf19 N-terminus as the receptor domain of the cohesin loading complex Scc2/4 (Hinshaw et al., 2017) enabled us to address whether Sli15/Ipl1 has an active role in the cohesin loading process. To this end, we deleted the first 30 amino acids of Ctf19 (Ctf19ΔN2-30) which contain the essential phosphorylation sites of the Dbf4-dependent kinase (DDK) required for the recruitment of the cohesin loading complex Scc2/4 to the centromere (Hinshaw et al., 2017). Strikingly, cells expressing Ctf19ΔN2-30 in the *sli15ΔN* background were indistinguishably viable upon depletion of Ctf19-FRB (Figure 5A), demonstrating that the synthetic lethality of *sli15ΔN/ctf19Δ* is independent of the Ctf19 N-terminus and its function in cohesin loading.

**Figure 5.**
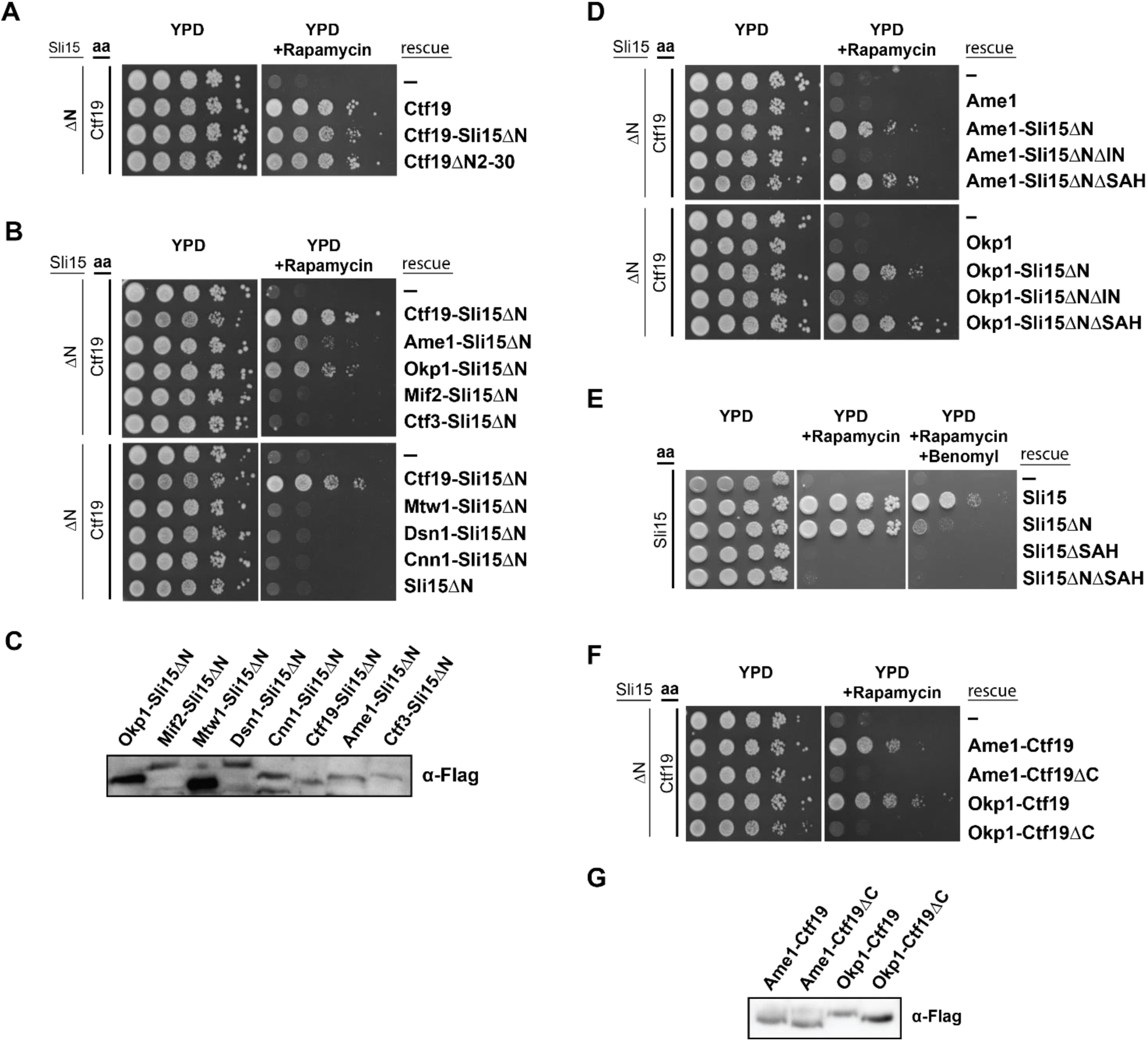
Synthetic lethality of centromere-targeting deficient Sli15 and Ctf19 depletion is rescued by fusing Sli15 exclusively to Ame1/Okp1 and is independent of Ctf19’s role in cohesin loading. (**A**) (**B**) (**D**) (**F**) Cell viability assays studying the rescue of the synthetically lethal *sli15ΔN/ctf19Δ* double mutant using the anchor-away system. The indicated constructs were transformed into a Ctf19 anchor-away (aa) strain (Ctf19-FRB) carrying *sli15∆N* (∆N) at the endogenous locus. Yeast growth was tested in serial dilutions either untransformed (-) or transformed with the indicated rescue constructs on YPD medium in the absence or presence of rapamycin at 30 °C. (**A**) Deletion of the Ctf19 N-terminus (Ctf19∆N2-30), required for cohesin loading, does not affect cell viability in a *sli15∆N* background. (**B**) Artificial and exclusive tethering of Sli15∆N to Ame1 or Okp1 rescued synthetic lethality of *sli15∆N* in Ctf19-FRB depleted cells. (**C**) Western blot analysis of the ectopically expressed protein levels of the inner and outer kinetochore subunits fused to Sli15∆N-7xFLAG in the yeast strains shown in (B). (**D**) Rescue of cell growth by exogenous Ame1-Sli15∆N or Okp1-Sli15∆N fusion proteins is dependent on the Sli15 Ipl1-binding domain (IN-box) whereas the SAH domain is dispensable. (**E**) Growth phenotypes of Sli15 wild-type, Sli15∆SAH, Sli15∆N, and Sli15∆N∆SAH tested in a Sli15-FRB anchor-away strain. Serial dilutions of Sli15-FRB strains ectopically expressing Sli15 wt and the indicated mutants were plated on YPD medium in the absence or presence of rapamycin or rapamycin and 15 μg/ml benomyl. Cells expressing Sli15∆N grow normally in the presence of rapamycin but exhibit benomyl sensitivity. Deletion of the SAH domain in Sli15 wt or Sli15∆N is lethal. (**F**) Growth assay analyzing the rescue of synthetic lethality of *sli15ΔN*/*Ctf19-FRB* strains by expressing Ame1-Ctf19, Okp1-Ctf19, Ame1-Ctf19∆C or Okp1-Ctf19∆C fusion proteins. (**G**) Western blot analysis visualizing the levels of exogenous C-terminally -7xFLAG tagged fusion proteins shown in (F).

To test whether synthetic lethality is associated with the loss of interaction between Sli15ΔN and COMA, artificial tethering of Sli15ΔN to the kinetochore should restore growth. We generated fusions of Sli15ΔN to various inner and outer kinetochore proteins and investigated whether growth was restored in a *Ctf19-FRB*/*sli15ΔN* background using the anchor-away system. Ectopic expression of Sli15ΔN fusions to the outer kinetochore subunits Mtw1 or Dsn1 and to the inner kinetochore subunits Mif2, Ctf3 or Cnn1 did not rescue viability (Figure 5B, C). However, selectively tethering Sli15ΔN to Ame1 or Okp1 restored growth (Figure 5B, C). Next, we tested whether the rescue of synthetic lethality was dependent on the Sli15 single alpha helix domain (SAH, aa 516-575) and the Ipl1 binding domain (IN-box, aa 626-698). Both domains are essential for cell growth in the *Sli15*wild-type or the *sli15ΔN* background (Figure 5E) (Kang et al., 2001). To discern the requirement for the two domains we generated Ame1-and Okp1-Sli15ΔN fusion constructs where either the IN-box (Adams et al., 2000, Kang et al., 2001), or the SAH domain (Samejima et al., 2015, van der Horst et al., 2015, Fink et al., 2017) of Sli15ΔN was deleted. While expression of Ame1- or Okp1-Sli15∆N∆SAH proteins rescued cell growth in the *sli15ΔN* background upon Ctf19 depletion, Ame1-and Okp1-Sli15∆N∆IN fusions did not, indicating that Ipl1 kinase activity is required (Figure 5D). Hence, since the ectopically expressed fusion proteins were tested in the *sli15ΔN* background, the result indicates that endogenous Sli15ΔN could not rescue *in trans* and that tethering Ipl1 activity exclusively to COMA subunits is crucial. In contrast, deletion of the SAH domain in Ame1- and Okp1-Sli15∆N∆SAH fusions was not lethal and could be rescued *in trans* (Figure 5D) suggesting that the SAH domain is not required for the function of CPC proximal to the centromere.

Since the RWD-C domain of Ctf19 was required for association with Sli15/Ipl1in our *in vitro* binding assay (Figure 4B), we asked whether this interaction is relevant for rescuing the Sli15ΔN/ΔCtf19 synthetic lethality. As recently described, the Ctf19 C-terminus is involved in formation of the COMA complex through binding to Okp1 (Schmitzberger et al., 2017), its deletion abrogates kinetochore localization of Ctf19/Mcm21. To circumvent loss of Ctf19/Mcm21 from kinetochores, we tested whether Ame1 or Okp1 fusions to wild-type Ctf19 or Ctf19∆C were able to rescue synthetic lethality in a *Ctf19-FRB/sli15∆N* background. Strikingly, fusions to wild-type Ctf19 restored viability whereas fusions to Ctf19∆C resulted in synthetic lethality (Figure 5F and 5G) indicating that positioning of Ipl1 activity proximal to Cse4 nucleosomes mediated by the Ctf19 C-terminus is important for CPC function.

## Discussion

### The Ame1/Okp1 heterodimer directly links Cse4 nucleosomes to the outer kinetochore

We investigated the subunit connectivity at the domain level to reveal the key differences in the inner kinetochore assembly at budding yeast point centromeres in comparison to regional centromeres in humans. Using *in vitro* reconstituted Cse4-NCPs and H3-NCPs in EMSAs, we screened stable subcomplexes of the budding yeast CCAN for their interaction with Cse4-NCPs. We discovered that in addition to Mif2, the Ame1/Okp1 heterodimer of the COMA complex is a direct and selective interactor of Cse4-NCPs (Figure 1A). To identify and characterize the binding interface between Ame1/Okp1 and Cse4 we applied crosslink-guided *in vitro* reconstitution where distance restraints and sequence conservation aided the detection of motifs for mutant analysis. This approach pinpointed the residues 163-187 of the Okp1 core domain (Figure 3B, C) and the residues 34-46 (Figure 2D, E) of a predicted helical motif within the Cse4 N-terminus as required for the selective Cse4 recognition. Sequence alignments of CENP-A orthologs from different species showed that the extended (130 aa) Cse4 N-terminus is unique to inter-related yeasts (Figure 2A) and identified the conserved END (aa 28 to 60) which is sufficient to provide the essential function of the Cse4 N-terminus (Chen et al., 2000). Furthermore, deletion of the N-terminal 50 amino acids of Cse4 was found to be lethal (Keith et al., 1999), whereas the H3 N-terminus was dispensable for viability (Mann and Grunstein, 1992), and END was previously implicated in interacting with the ‘Mcm21/Okp1/Ctf19’ complex (Ortiz et al., 1999, Chen et al., 2000). Consistent with these previous findings, we narrowed down the essential N-terminal Cse4 patch to amino acids 34-46 which was identified as the minimal motif sufficient for the interaction with Ame1/Okp1. The notion that the essential function of the Cse4 N-terminus and the binding interface for Ame1/Okp1 are mediated by the same 13 amino acid long motif suggests that the interaction of the Ame1/Okp1 complex with Cse4 is essential in budding yeast.

The Ame1/Okp1 heterodimer in complex with the Cse4-NCP retained the ability to associate with the outer kinetochore MTW1c (Figure 2C) (Hornung et al., 2014). As MTW1c recruitment by Mif2 and Cnn1 are redundant pathways and become only essential if one of both is compromised (Hornung et al., 2014, Lang et al., 2018), Ame1/Okp1 is the sole essential link of the centromeric nucleosomes to the outer kinetochore which emphasizes its role as cornerstone of kinetochore assembly at the budding yeast point centromere.

### Dual recognition of Cse4 at point centromeres by a CCAN architecture distinct from vertebrate regional centromeres

In vertebrates two CCAN proteins, CENP-N and CENP-C, were shown to directly and selectively interact with CENP-A. CENP-C binds divergent hydrophobic residues of the CENP-A C-terminal tail, whereas CENP-N associates with the CENP-A CATD (Carroll et al., 2009, Carroll et al., 2010, Guse et al., 2011, Kato et al., 2013, Weir et al., 2016, Pentakota et al., 2017). Up to now, only Mif2 has been found to bind the C-terminus of Cse4 (Xiao et al., 2017). Interestingly, our EMSA and XLMS data suggest that in contrast to the human ortholog, CENP-N/L, the Chl4/Iml3 heterodimer does not directly interact with Cse4-NCP, but is linked to Ame1/Okp1 and Mif2 (Figure 1A, C). Recent electron microscopy reconstructions of human CENP-A nucleosomes in complex with CENP-N/L identified the RG loop in the CATD (Zhou et al., 2011) to be required for CENP-N interacting with CENP-A (Chittori et al., 2018, Pentakota et al., 2017, Tian et al., 2018). The lack of this RG loop in the budding yeast Cse4 CATD is consistent with our finding of Chl4/Iml3 being positioned distal from Cse4 nucleosomes (Figure 1A, C). In humans, recruitment of the CENP-OPQRU complex to kinetochores requires a joint interface formed by CENP-HIKM and CENP-LN (Okada et al., 2006, Foltz et al., 2006, Pesenti et al., 2018) and loss of the complex does not affect localization of other inner kinetochore proteins. The distinct architectures of vertebrate and budding yeast inner kinetochores are reflected by the physiological importance of the involved proteins, as Ame1/Okp1 together with Mif2 are the essential CCAN proteins in budding yeast whereas knockouts of CENP-U/Q in DT40 cells are viable (Hori et al., 2008). Our *in vitro* reconstitution study of the budding yeast inner kinetochore unveils the molecular foundation of the essential function of the Cse4 N-terminus and demonstrates the key role of Ame1/Okp1 in establishing the Cse4 selective binding environment in budding yeast. The fact that both Cse4 interactors, Mif2 and Ame1/Okp1, directly link to the outer kinetochore MTW1c could reflect the requirement for recruiting additional KMN complexes through the individual Ame1 and Mif2 N-termini (Przewloka et al., 2011, Screpanti et al., 2011, Hornung et al., 2014) or for building a high-affinity binding site for the Cse4-NCP through cooperativity which both would facilitate force transmission across a single kinetochore unit upon microtubule attachment. These architectural differences between budding yeast and human CCAN complexes may be the consequence of building a kinetochore unit at a point centromere linked to a single microtubule in budding yeast, in contrast to vertebrate regional centromeres, where an array of those kinetochore units provide attachment sites for 3 to 30 microtubules (Walczak et al., 2010).

### COMA and its role in positioning Sli15/Ipl1 at the inner kinetochore

Besides the two essential subunits Ame1/Okp1 of the COMA complex the Ctf19/Mcm21 heterodimer is not essential. *Ctf19∆* and *mcm21∆* mutants are viable but exhibit chromosome segregation and cohesion defects (Ortiz et al., 1999, Hyland et al., 1999, Poddar et al., 1999, Fernius and Marston, 2009, Hinshaw et al., 2017). Interestingly, Ctf19 or Mcm21 become essential for viability once Sli15 loses its ability to be recruited to the inner centromere by deleting the Sli15 N-terminus (Sli15∆N). This observation has led to the hypothesis that centromere-anchored Sli15 might be involved in the cohesin loading or maintenance process (Campbell and Desai, 2013). An alternative model posits that COMA is required for the precise positioning of Sli15/Ipl1 at the kinetochore and the importance of this function is only revealed if centromere targeting is perturbed. Supporting this scenario a role of COMA in the localisation of Sli15 at kinetochores and the regulation of CPC function has been proposed earlier (Knockleby and Vogel, 2009).

Our crosslink-guided *in vitro* reconstitution and binding assays revealed that COMA directly interacts with Sli15/Ipl1 and pinpointed the C-terminal RWD domain of Ctf19 as primary docking site (Figure 4B). Importantly, synthetic lethality upon Ctf19 or Mcm21 depletion in the *sli15∆N* mutant background was exclusively rescued by fusions of Sli15∆N to COMA subunits whereas other inner or outer kinetochore protein fusions failed to rescue growth (Figure 5B). This observation indicates that the positioning of Sli15/Ipl1 proximal to COMA or even their direct interaction is important *in vivo*. Because of the requirement of a functional Ipl1 binding IN-box of Sli15 for restoring viability we assume that the observed synthetic lethality is due to mislocalized Ipl1 kinase (Figure 5D). Tethering Sli15 to the inner kinetochore might ensure the precise spatial positioning of Ipl1 kinase towards outer kinetochore substrates like Ndc80, Dam1 or Dsn1 (Foley and Kapoor, 2013, Akiyoshi et al., 2013, Krenn and Musacchio, 2015) required to detect and destabilize erroneous microtubule attachments or to stabilize outer kinetochore recruitment, respectively (Figure 6). In contrast, COMA-Sli15ΔN fusions lacking the SAH domain were able to rescue growth indicating that this domain is dispensable for CPC function at the inner kinetochore. However, since the SAH domain is required for binding to spindle microtubules and critical for cell survival (Samejima et al., 2015, van der Horst et al., 2015, Fink et al., 2017), we conclude that the observed rescue was mediated *in trans* by Sli15ΔN (Figure 5D, E). Strikingly, we were able to demonstrate that Sli15ΔN was not synthetically lethal with Ctf19ΔN2-30 lacking the Ctf19 N-terminus which was identified as receptor for the cohesin loading complex (Hinshaw et al., 2017). This pointed towards a mechanism involving another domain of Ctf19. On the basis of crosslink-derived restraints and *in vitro* reconstitution experiments, we showed that deletion of the C-terminal RWD domain of Ctf19 (Ctf19∆C) is sufficient to cause synthetic lethality with Sli15ΔN (Figure 5F) and that recombinant Ctf19∆C in complex with Mcm21 (Figure 4B) abrogated interaction with Sli15. Taken together, the *in vitro* and *in vivo* data clearly indicate that the C-terminus of Ctf19 acts as a primary Sli15 binding site within COMA, suggesting that Ctf19/Mcm21, linked to Cse4-NCPs through Ame1/Okp1, is important for positioning Ipl1 activity close to Cse4 nucleosomes in a Bir1 independent manner (Jeyaprakash et al., 2007, Cho and Harrison, 2011). We demonstrated that the interaction of Sli15 with Ctf19 is functionally important in budding yeast suggesting that positioning of Ipl1 at the inner kinetochore may be part of a distinct kinetochore conformation that facilitates the selective recognition and destabilization of erroneous microtubule attachments upon loss of tension. Whether the observed interaction between the CPC and COMA is conserved in higher eukaryotes or whether this interaction is facilitated by other kinetochore proteins is unclear. Interestingly, the orthologous human CENP-OPQUR complex has recently been shown to promote accurate chromosome alignment by interaction with microtubules (Pesenti et al., 2018).

**Figure 6.**
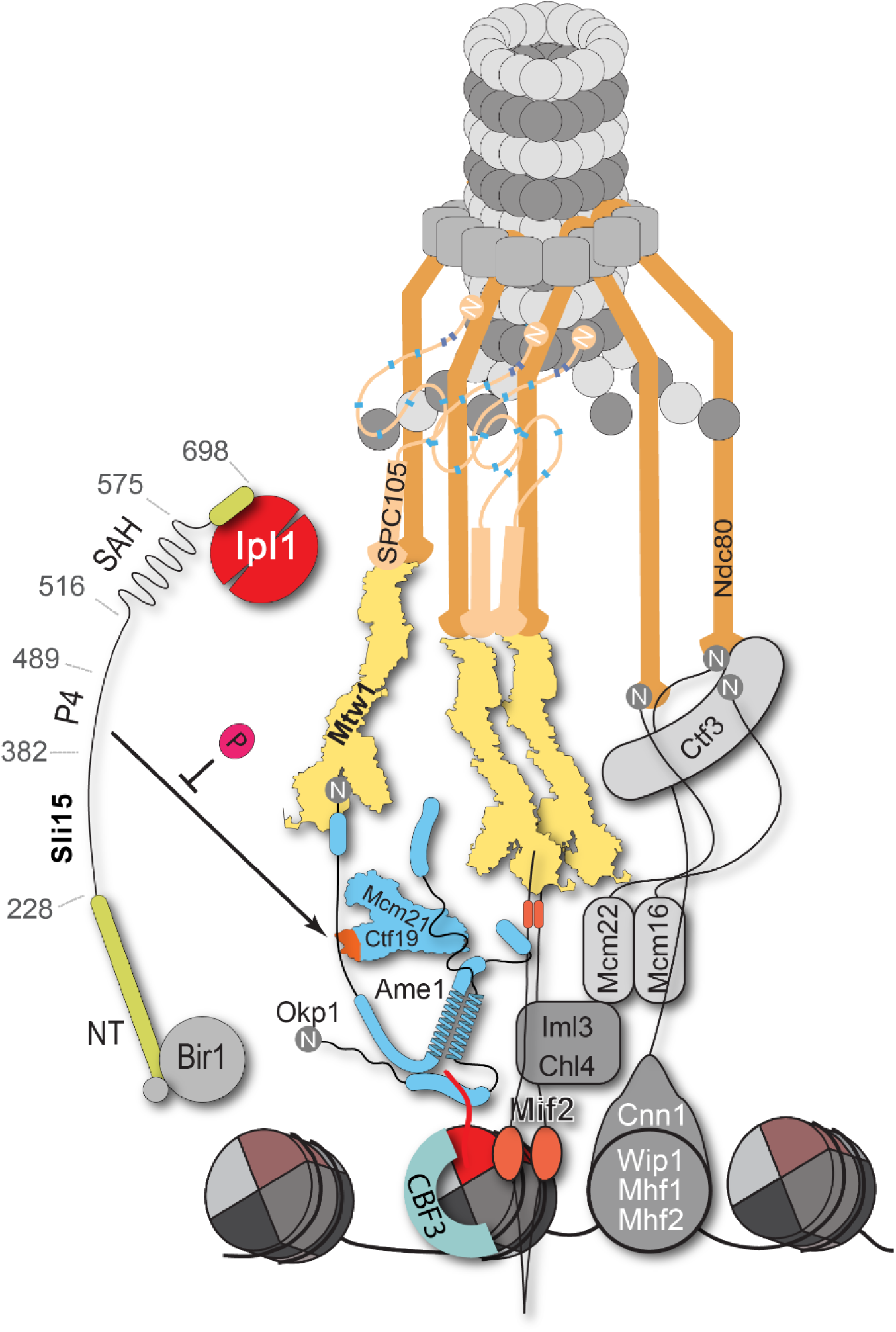
Schematic model of the budding yeast kinetochore subunit architecture and its interaction with Sli15/Ipl1. Our finding that the Okp1 core domain directly binds the essential motif of the Cse4 END domain, whereas Chl4 does not recognize Cse4-NCPs, suggests that in contrast to humans, the dual recognition of Cse4-NCPs in *S. cerevisiae* is established by Ame1/Okp1 and Mif2. Ame1/Okp1 is thus central to kinetochore assembly in budding yeast and together with the observation that Ctf19 associates with Sli15/Ipl1, the COMA complex may contribute to position Ipl1 kinase close to centromeric chromatin in order to facilitate feedback control of chromosome segregation.

Here, we used the high-performance XLMS technique to identify so far unknown protein-protein interactions and characterized the physiological relevance of their interfaces *in vivo*. Our observations support a model that unveils COMA as the centrepiece of kinetochore assembly in budding yeast (Figure 6) and contributes to the molecular understanding of the fascinating question of how cells establish correct chromosome biorientation at the mitotic spindle.

## Materials and methods

### Chemical crosslinking and mass spectrometry of kinetochore complexes

The complex containing Cse4-NCP, Mif2, Ame1/Okp1, Ctf19/Mcm21, Chl4/Iml3 and MTW1c was assembled in solution. It was cross linked using an equimolar mixture of isotopically light (hydrogen) and heavy (deuterium) labeled bis[sulfosuccinimidyl]suberate (BS3, H12/D12) (Creative Molecules) at a final concentration of 0.25 - 0.5 mM at 10 °C for 30 min. The reaction was quenched by adding ammonium bicarbonate to a final concentration of 100 mM for 10 min at 10 °C. The sample was submitted to size exclusion chromatography on a Superose 6 Increase 10/300 GL column (GE Healthcare) and the fractions corresponding to the crosslinked complex were selected for the subsequent protein digest and mass spectrometry (described further down). The complex of Sli15-2xStrep-HA-6xHis/Ipl1 with Ame1/Okp1 and Ctf19/Mcm21 was assembled on Strep-Tactin Superflow agarose (Qiagen) by incubation at room temperature (RT), 1000 rpm for 1 h in a thermomixer (Eppendorf). Unbound proteins were removed by washing three times with binding buffer (50 mM NaH_2_PO_4_ pH 8, 500 mM NaCl, 5% glycerol) and the complex was eluted in binding buffer containing 8 mM biotin. The eluted complex was re-isolated on Ni-NTA (Qiagen) beads, washed twice with binding buffer and then crosslinked by resuspending the protein bound beads in BS3 cross linker at a final concentration of 0.25-0.5 mM at 30 °C for 30 min. The crosslinking reaction was stopped by adding ammonium bicarbonate to a final concentration of 100 mM for 20 min at 30 °C.

Crosslinked samples were denatured by adding 2 sample volumes of 8 M urea, reduced with 5 mM TCEP (Thermo Scientific) and alkylated by the addition of 10 mM iodoacetamide (Sigma-Aldrich) for 40 min at RT in the dark. Proteins were digested with Lys-C (1:50 (w/w), Wako Pure Chemical Industries) at 35 °C for 2 h, diluted with 50 mM ammonium bicarbonate, and digested with trypsin (1:50 w/w, Promega) overnight. Peptides were acidified with trifluoroacetic acid (TFA) at a final concentration of 1% and purified by reversed phase chromatography using C18 cartridges (Sep-Pak, Waters). Crosslinked peptides were enriched on a Superdex Peptide PC 3.2/30 column using water/acetonitrile/TFA (75/25/0.1, v/v/v) as mobile phase at a flow rate of 50 μl/min and were analyzed by liquid chromatography coupled to tandem mass spectrometry (LC-MS/MS) using an Orbitrap Elite instrument (Thermo Fisher Scientific). Fragment ion spectra were searched and crosslinks were identified by the dedicated software *xQuest* (Walzthoeni et al., 2012). The results were filtered according to the following parameters: Δ score ≤ 0.85, MS1 tolerance window of -4 to 4 ppm and score ≥ 22. The quality of all crosslink spectra passing the filter was manually validated and crosslinks were visualized as network plots using the webserver *xVis* (Grimm et al., 2015).

### Electrophoretic mobility shift assay

Reconstituted nucleosomes and protein complexes were mixed in a 1:1 or 1:2 molar ratio in a buffer containing 20 mM Hepes (pH 7.5) and incubated for 1h on ice. The interaction was analyzed by electrophoresis at 130 V for 70–90 min on a 6% native polyacrylamide gel in a buffer containing 25 mM Tris and 25 mM boric acid. After electrophoresis, gels were stained with SYBR^®^ Gold (Thermo Fisher).

### Analytical size exclusion chromatography for interaction studies

Analytical SEC experiments were performed on a Superdex 200 Increase 3.2/300 column (GE Healthcare). To detect the formation of a complex, proteins were mixed at equimolar ratios and incubated for 1 h on ice before SEC. All samples were eluted under isocratic conditions at 4 °C in SEC buffer (50 mM HEPES pH 7.5, 150 mM NaCl, 5% glycerol). Elution of proteins was monitored by absorbance at 280 nm. 100 μl fractions were collected and analyzed by SDS-PAGE and Coomassie staining.

### *In vitro* protein binding assay of Sli15/Ipl1 to Ame1/Okp1 and/or Ctf19/Mcm21

Phosphorylated or non-phosphorylated wild-type or mutant Sli15-2xStrep-HA-6xHis/Ipl1 was immobilized on Strep-Tactin Superflow agarose (Qiagen). For prephosphorylation, Sli15/Ipl1 was incubated at 25 °C for 25 min in the presence of 3 mM MgCl_2_ and 3 mM ATP. Samples for non-phosphorylated Sli15/Ipl1 were treated the same way but instead of 3 mM ATP the non-hydrolysable analog AMP-PNP (Santa Cruz Biotechnology) was applied. Subsequently, non-phosphorylated as well as phosphorylated Sli15/Ipl1 complexes were washed three times with binding buffer (50 mM NaH_2_PO_4_ pH 8, 120 mM NaCl, 5% glycerol).

Testing of binding between Ame1/Okp1, Ctf19/Mcm21 and Sli15/Ipl1 was performed in binding buffer at 4 °C, 1000 rpm for 1 h in a thermomixer (Eppendorf). Unbound proteins were removed by washing three times with binding buffer. The complexes were either eluted with 8 mM biotin in 50 mM NaH_2_PO_4_ pH 8, 500 mM NaCl, 5% glycerol, or by boiling in 2x SDS loading buffer.

### Amino acid sequence alignment

Multiple sequence alignment of Cse4 or Okp1 protein sequences from interrelated budding yeast species was conducted with Clustal Omega (Sievers et al., 2011). Only protein sequences with the highest similarity to *S. cerevisiae* Cse4 or *S. cerevisiae* Okp1 as determined by a protein BLAST search were included in the search. In addition three mammalian and the *Schizosaccharomyces pombe* homologous CENP-A protein sequences were included in the Cse4 alignment.

### Structural model prediction of Cse4

The model was predicted using I-Tasser (Yang et al., 2015) upon addition of 6 intramolecular contact restraints, which were observed in a XLMS study of the Cse4-nucleosome with inner kinetochore proteins (Figure 1C). The model was aligned to histone H3 (HHT1) from the crystallized *S. cerevisiae* nucleosome core particle (PDB entry 1ID3). The alignment was performed in Chimera (Pettersen et al., 2004) applying the match maker tool, following alignment pair of chains and the Needleman-Wunsch algorithm (Needleman and Wunsch, 1970).

### Yeast strains and methods

All plasmids and yeast strains used in this study are listed in tables S1 and S2, respectively. Yeast strains were created in the S288C background. The generation of yeast strains and yeast methods were performed by standard procedures. The anchor-away technique was performed as previously described (Haruki et al., 2008).

For anchor-away rescue experiments, the respective promoters and coding sequences were PCR amplified from yeast genomic DNA and cloned into the vector pRS313 either via the Gibson assembly or the restriction/ligation method. In order to artificially target Sli15∆N2-228 to the kinetochore, the individual promoters and genes were PCR amplified and the respective gene fusions [*CTF19*, *AME1*, *OKP1*, *CTF3*, *CNN1*, *MIF2*, *DSN1*, *MTW1*]-[*SLI15∆N2-228*]-[*6xHis-7xFLAG*] (table S1) were generated and cloned into pRS313 using the Gibson assembly reaction The individual deletion mutants were generated using the Q5 site-directed mutagenesis kit (New England Biolabs). The rescue constructs were transformed into Cse4-, Ctf19-, Okp1-, or Sli15 anchor-away strains and cell growth was tested in 1:10 serial dilutions on YPD plates in the absence or presence of rapamycin at 30 °C for 72 hours.

### Western blot analysis

For western blot analysis an equivalent of 10 OD_600_ of cells logarithmically grown in liquid culture was collected by centrifugation at 3140 x g for 5 min at RT and the pellet was washed once with aqua dest. For protein extraction, the pellet was resuspended in 1 ml ice-cold 10% trichloroacetic acid and incubated on ice for 1h. Samples were pelleted at 20000x g for 10 min, 4 °C and washed twice with ice-cold 95% ethanol. Pellets were air-dried and resuspended in 100 μl 1x SDS-PAGE sample buffer containing 75 mM Tris pH 8.8. Samples were boiled (10 min 95 °C) and centrifuged at 10800 x g for 3 min at RT and supernatants were separated on 10% SDS-PAGE gels. Immunoblotting was performed with the following antibodies: Anti-FLAG M2 (Sigma-Aldrich) and visualized by HRP-conjugated anti-mouse secondary antibodies (Santa Cruz).

### Protein expression and purification

Expression constructs for 6xHis-Chl4/Iml3, 6xHis-Cnn1/Wip1-1xFlag, 6xHis-Nkp1/Nkp2 and Mhf1/Mhf2-1xStrep were created by amplification of genomic DNA and cloned into pETDuet-1 vector (Novagen). Expression was performed in BL21 (DE3) cells (New England Biolabs). Cells were grown at 37 °C until OD_600_ 0.6, followed by induction with 0.5 mM IPTG for Chl4/Iml3 or 0.2 mM IPTG for all other protein expressions. Protein expression was induced overnight at 18 °C, or for 3h at 23 °C, respectively.

Cells were lysed using a French Press in lysis buffer (50 mM Hepes pH 7.5, 400 mM NaCl, 3% glycerol, 0.01% Tween20 and protease inhibitor cocktail (Roche). 6xHis-tagged proteins were purified using Ni-NTA agarose (Qiagen), whereby 30 mM imidazole were added to the lysis buffer in the washing step, followed by protein elution in 50 mM Hepes pH 7.5, 150 mM NaCl, 300 mM imidazole, and 5% glycerol. Strep-tag purification was performed using Strep-Tactin Superflow agarose (Qiagen) and eluted in a buffer containing 50 mM Hepes pH 7.5, 150 mM NaCl, 8 mM biotin, and 5% glycerol.

Buffer exchange into a buffer containing 50 mM Hepes pH 7.5, 150 mM NaCl, and 5% glycerol was performed using a Superdex 200 HiLoad 16/60 column (GE-Healthcare) for Chl4/Iml3 and Cnn1/Wip1 or using a PD10 desalting column (GE-Healthcare) for Nkp1/2 and Mhf1/2 protein complexes.

### Ame1/Okp1 expression and purification

Ame1-6xHis/Okp1 wild-type and mutant protein expression and purification in *E. coli* was performed as described previously (Hornung et al., 2014).

### *In vitro* reconstitution of Cse4- and H3-NCPs

Octameric Cse4 and H3 containing nucleosomes were *in vitro* reconstituted from budding yeast histones which were recombinantly expressed in *E.coli* BL21 (DE3) and assembled on 167 bp of the ‘Widom 601’ nucleosome positioning sequence according to a modified protocol of (Turco et al., 2015).

### Affinity-purification of recombinant protein complexes from insect cells

C-terminal 6xHis-6xFLAG-tags on Mcm21, Mif2, Dsn1, Mcm16 and C-terminal 2xStrep-tags on Sli15 were used to affinity-purify Ctf19/Mcm21, Mif2, MTW1c, CTF3c and Sli15/Ipl1 complexes. Open reading frames encoding the respective subunits were amplified from yeast genomic DNA and cloned into the pBIG1/2 vectors according to the biGBac system (Weissmann et al., 2016). The pBIG1/2 constructs were used to generate recombinant baculoviral genomes by Tn7 transposition into the DH10Bac *E. coli* strain (Thermofisher) (Vijayachandran et al., 2011). Viruses were generated by transfection of Sf21 insect cells (Thermo Scientific) with the recombinant baculoviral genome using FuGENE HD transfection reagent (Promega). Viruses were amplified by adding transfection supernatant to Sf21 suspension cultures. Protein complexes were expressed in High Five insect cell (Thermo Scientific) suspension cultures.

For purification of FLAG-tagged kinetochore complexes, insect cells were extracted in lysis buffer (50 mM Tris pH 7.5, 150 mM NaCl, 5% glycerol) supplemented with Complete EDTA-free protease inhibitors (Roche) using a Dounce homogenizer. Cleared extracts were incubated with M2 anti-FLAG agarose (Sigma-Aldrich) for 2 h, washed three times with lysis buffer and eluted in lysis buffer containing 1 mg/ml 3xFLAG peptide.

High Five cells expressing Strep-tagged Sli15/Ipl1 were lysed in 50 mM NaH_2_PO_4_ pH 8, 300 mM NaCl, 5% glycerol supplemented with Complete EDTA-free protease inhibitors (Roche). Subsequent to incubating the cleared lysates with Strep-Tactin Superflow agarose (Qiagen), protein bound beads were washed three times with lysis buffer and the bound protein complex was eluted in lysis buffer containing 8 mM biotin. FLAG peptide or biotin was either removed via PD10 desalting columns (GE-Healthcare) or SEC using a Superdex 200 HiLoad 16/60 column (GE-Healthcare) and isocratic elution in lysis buffer.

## Acknowledgements

We are grateful to Andrea Musacchio (MPI Dortmund) and Stefan Westermann (University of Essen, Germany) for discussions and sharing reagents. We thank Alwin Köhler (Max F. Perutz Laboratories, Vienna) for sharing the protocol for nucleosome reconstitution. JFH and GH were funded by the Graduate School (GRK 1721) and MP and VS were funded by the Graduate School (Quantitative Biosciences Munich) of the German Research Foundation (DFG). FH was supported by the European Research Council (ERC-StG no. 638218), the Human Frontier Science Program (RGP0008/2015), by the Bavarian Research Center of Molecular Biosystems and by a LMU excellent junior grant.

## Supplementary figures

**Figure S1.**
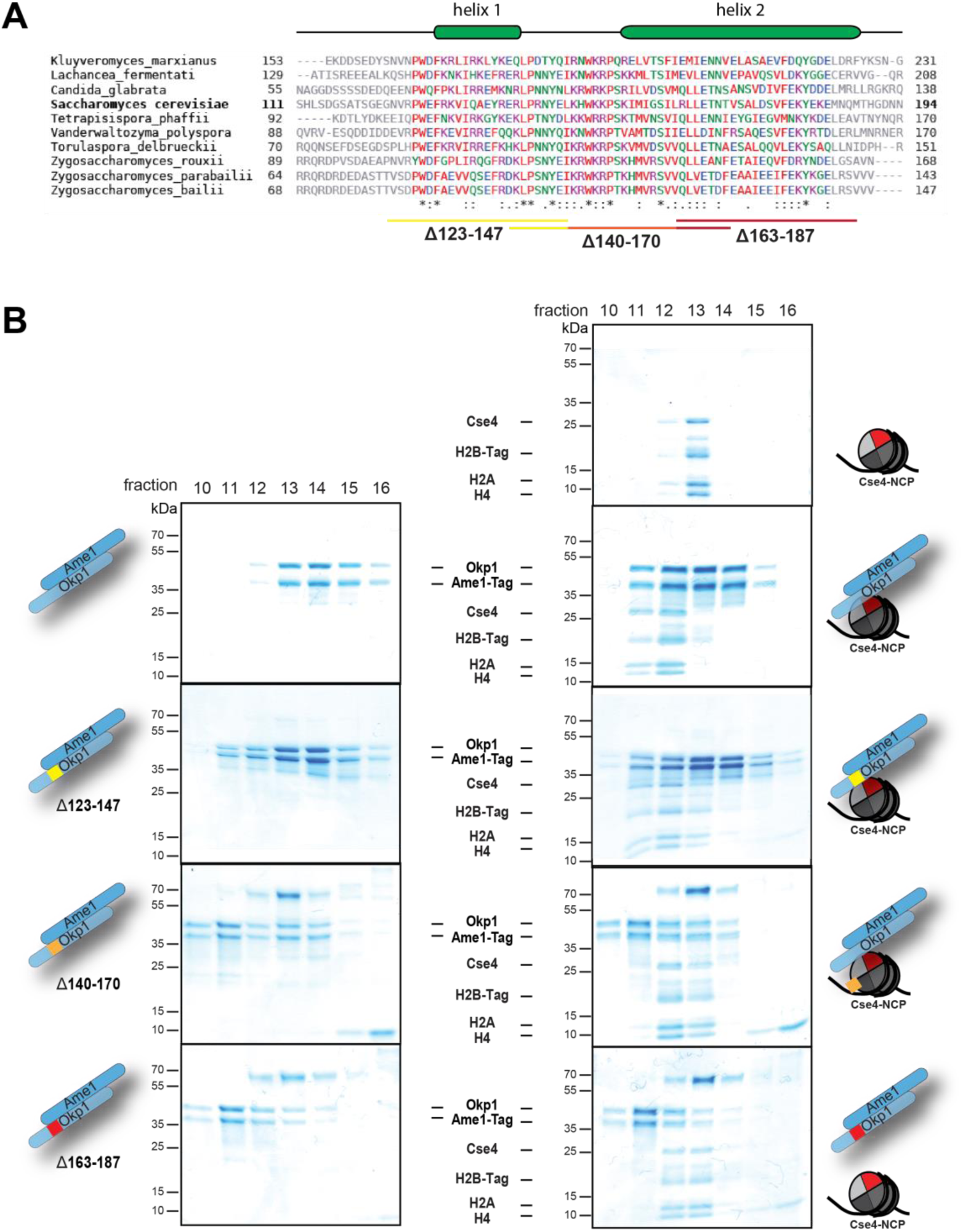
Identification of the Cse4 binding site on Okp1. **(A)** Multiple sequence alignment of Okp1 amino acid sequences from related yeast species. Amino acid residues of the conserved region are colored and annotated according to the ClustalW color and annotation codes. Green bars above the alignment represent alpha helical regions predicted by Jpred (Drozdetskiy et al., 2015). Lines below the alignment indicate the overlapping Okp1 deletion mutants analysed in (B). **(B)** Size exclusion chromatography (SEC) analysis of equimolar mixtures of reconstituted Cse4-NCPs with recombinant wild-type Ame1/Okp1, Ame1/Okp1Δ123-147, Ame1/Okp1Δ140-170 or Ame1/Okp1Δ163-187 mutant complexes. Eluted proteins were visualized by SDS-PAGE and Coomassie staining.

**Figure S2.**
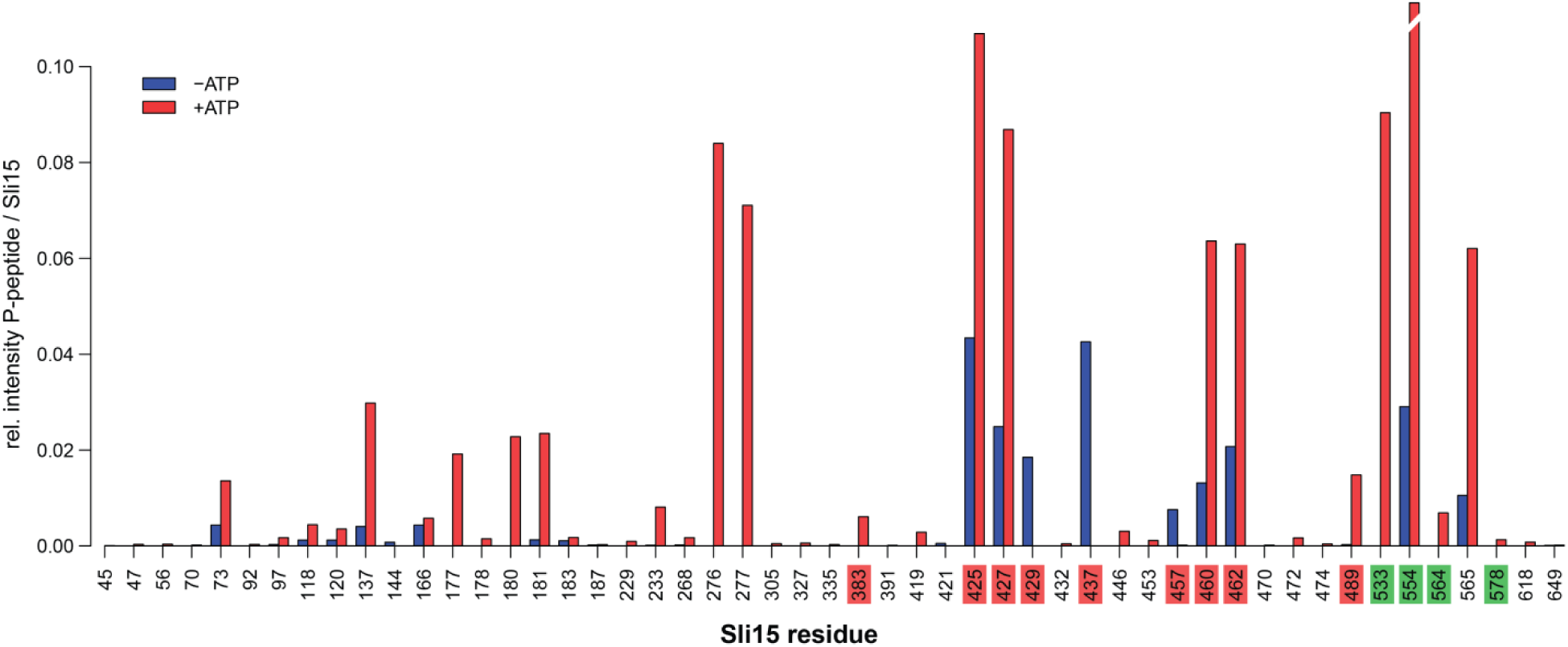
Quantification of Ipl1 *in vitro* phosphorylation sites on Sli15 represented as barplot. The Sli15/Ipl1 complex was prepared in untreated (blue bars) and auto-phosphorylated (red bars) states and phosphorylation sites were quantified by mass spectrometry and the software MaxQuant (vers. 1.5.2.8) (Cox and Mann, 2008). Protein abundances were estimated as the sum of peptide intensities divided by the peptide counts. The intensities of the phosphorylated peptides were normalized to the respective protein intensities. Barplots represent the average of three replicates per condition with non-assigned values removed prior to averaging. Phosphorylation sites of Sli15 highlighted in red or green were mutated to aspartate resulting in the phosphorylation-mimicking Sli15 P4 (red) or P7 (green) mutants which are also depicted in the scheme shown in Figure 4C.

